# Canopies drive the reassembly of pollinator communities and interaction networks along tropical forest succession

**DOI:** 10.1101/2025.01.23.634515

**Authors:** Ugo Mendes Diniz, Dennis Böttger, Sabine Fernanda Viteri Lalama, Kilian Frühholz, Maximilian Pitz, Julia Windl, Claus Rasmussen, Alexander Keller, Sara Diana Leonhardt, Gunnar Brehm

## Abstract

Vertical stratification is a prominent driver of biodiversity in forests. As the proportion of successional forests increases worldwide and canopies are lost at a fast pace, it is crucial to understand the role of stratification as a driver of community recovery and ecological processes during succession, particularly for important plant mutualist groups essential for forest recovery. Within a well-resolved chronosequence in the northwestern Ecuadorian rainforest, we compiled an extensive database of over 20,000 diurnal and nocturnal pollinators and 2,000 interactions with plants and examined the interacting effects of recovery age and stratification on pollinator community and interaction network reassembly. In late successional and old-growth forests, stratification was a stronger predictor of pollinator abundance, alpha-diversity, functional richness and beta-diversity than forest legacy. While most groups were strongly associated with canopies (moths, social bees, and nocturnal bees), orchid bees exhibited an inverse pattern, also presenting the strongest functional response to stratification and recovery status. Within the entire chronosequence (0-38 years of recovery, plus old-growth forests), recovery status and stratification, alongside landscape features, acted synergistically as predictors of pollinator community parameters. Although structurally stable across the chronosequence and between strata, Interaction networks were richer and most distinct in canopies, while the highest pollinator and interaction diversity were found in old-growth canopies. In successional forests, networks were only comparable in size to active disturbances and early recovery with both strata combined, whereas complete networks in old-growth forests were more than twice as large as those in other recovery statuses. Our study underlines the importance of stratification in leveraging pollinator diversity during forest succession. By serving as havens for canopy-adapted species, tropical canopies may thus help safeguard pollination processes in disturbed landscapes.

## 1. INTRODUCTION

Forests are remarkably diverse ecosystems that harbor most of the Earth’s biota [^1^]. The diversity of biological niches offered by forests can ultimately be linked to their multidimensional complexity in the form of habitat heterogeneity (clearings, edges, and patch-matrix dynamics) [^2^] and a wide range of microhabitats and resources associated with trees, epiphytes, and other midstory components [^3^]. One of the most prominent drivers of forest heterogeneity and biotic composition is vertical stratification, characterized by distinct niches formed through the interplay of microclimatic conditions, microhabitats, and resources [^4,5^].

While the forest understory offer a variety of resources associated with the soil (e.g., dung, deadwood, carrion, and soil microfauna) and with low-laying vegetation (herbs and bushes in deciduous forests or pioneers in clearings) [^6^], resources essential to certain consumer guilds are concentrated in the forest’s upper layer, particularly in tropical rainforests where a clear degree of vertical stratification is present [^6^]. The canopy, a photosynthetic hotspot in the forest due to increased radiation, excels in resource offer for primary consumers [^7^]. Large and productive trees have a much higher resource yield (e.g., flowers and fruits) than understory plants [^8^], while the accumulation of epiphytes, lianas, and tree crown results in a higher taxonomical and functional diversity of primary resources and turns canopies into optimal foraging areas for herbivores, frugivores and nectarivores [^9,10^]. Indeed, a quarter of all invertebrates are expected to be canopy specialists [^6^].

Nonetheless, canopies are fragile and affected not only by acute forest clearings but also by the chronic targeted logging of slow-growing, large hardwood trees that comprise a significant portion of canopies [^11^]. Furthermore, canopies suffer from an inherently low resilience, as veteran trees may take centuries to establish, grow, and provide their ecosystem functions [^12^]. Tropical forests are particularly critical in this regard. They foster an estimated three-quarters of Earth’s biodiversity and account for most of the world’s current Biodiversity Hotspots [^13^]. In these ecosystems, the loss of vertical complexity through selective logging, combined with forest fragmentation and reduction in patch size, significantly reduces the incidence of essential low-mobility canopy specialists, such as fruit-dispersing primates [^5^]. However, these stressors also filter out a relevant proportion of important starters and maintainers of forest succession and recovery from the landscape, such as mobile canopy-specialist pollinators and seed dispersers [^14,15^]. Even partial canopy losses may lead to significantly higher exposure in forest understories due to increased heat and light incidence, which further affects species occurrences in forests affected by targeted logging, especially among desiccation-prone animals such as many arthropod groups [^6^].

Therefore, it is crucial to understand the impact of canopy loss on arthropod groups that partake in essential ecosystem processes necessary for forest recovery. Insect pollinators, for instance, are critical for the regeneration of tropical forests [^16^]. The consequences of losing canopy-associated habitats, however, are still relatively little understood for insects. Most of our current knowledge of invertebrate diversity is still based on understory sampling [^17^], which stems from logistical limitations in sampling canopies, often leading to small sample sizes and replication issues [^17,18^]. While there is evidence that several insect communities, including pollinators, may be more diverse in canopies [^19–21^], data are sparse and often conflicting, and the ecological drivers of pollinator stratification are understudied [^17,22–25^]. Pollinators may be particularly associated with the presence of resource-dense canopy trees in primary or old successional forests, as they provide abundant and year-round resources for several groups of flower-visiting insects [^9,20,26^]. The role of canopy formation as a driver of pollinator communities and the pollination service they provide during the process of forest succession has, to date, not been explicitly investigated, thereby precluding our understanding of the reassembly of plant-pollination networks.

Therefore, we examined whether stratification significantly affects pollinator communities and their interaction networks with plants along a tropical forest regeneration gradient. Ultimately, we sought to understand the role of canopy formation on the recovery of a key mutualistic ecosystem process. Specifically, we expected:

i. Stratification to enhance pollinator diversity along forest recovery by supplying a distinct environment in successional forests that can be colonized by canopy-adapted species. Therefore, canopies shall yield communities complementary to understories, leading to higher overall abundance and (functional) diversity in canopy-bearing forests. This effect will be evident after the first few decades of recovery but will be stronger in older forests, where floral resource stratification is more pronounced.
ii. Pollinator community complementarity to be driven by pollinators’ interaction, dispersal, and response traits. Given the different and presumably richer assemblage of flowering species in the canopy, we expect interaction traits (e.g., proboscis length) to be more variable in canopies. Accordingly, dispersal traits in canopy-loving insect assemblages (e.g., larger body size and high flight efficiency), as well as response traits (e.g., incidence of aposematism and crypticism), will reflect the drier, warmer, and windier conditions found in canopies, as well as an increased exposure to predators.
iii. Interaction networks will be larger and more diverse in canopies due to larger pools of pollinator and plant species interacting, and present the most distinct interactions due to being composed of canopy-specialist species. Network size will be considerably larger in old-growth forests due to increased community divergence between strata, resulting in higher complementarity.

## METHODS

### Study site

The study was conducted in 2022 and 2023 in the Río Canandé (0°31’33.4“N, 79°12’46.0” W) and the Tesoro Escondido (0°32’30.9“N, 79°08’41.9”W) Reserves, Esmeraldas province, northwestern Ecuador (Fig. 1A). The site is a lowland forest ecosystem within the Chocó-Darien ecoregion, with a mean annual temperature of 24°C and annual rainfall of ca. 2100 mm, (Af climate, Köppen scale). The study was carried out in an ~7800 ha area within the REASSEMBLY research unit, a well-resolved chronosequence of 62 50×50 m plots ranging from active disturbance agricultural sites to regenerating secondary forests and old-growth forests [^27^] (Fig. 1B).

**Figure 1.**
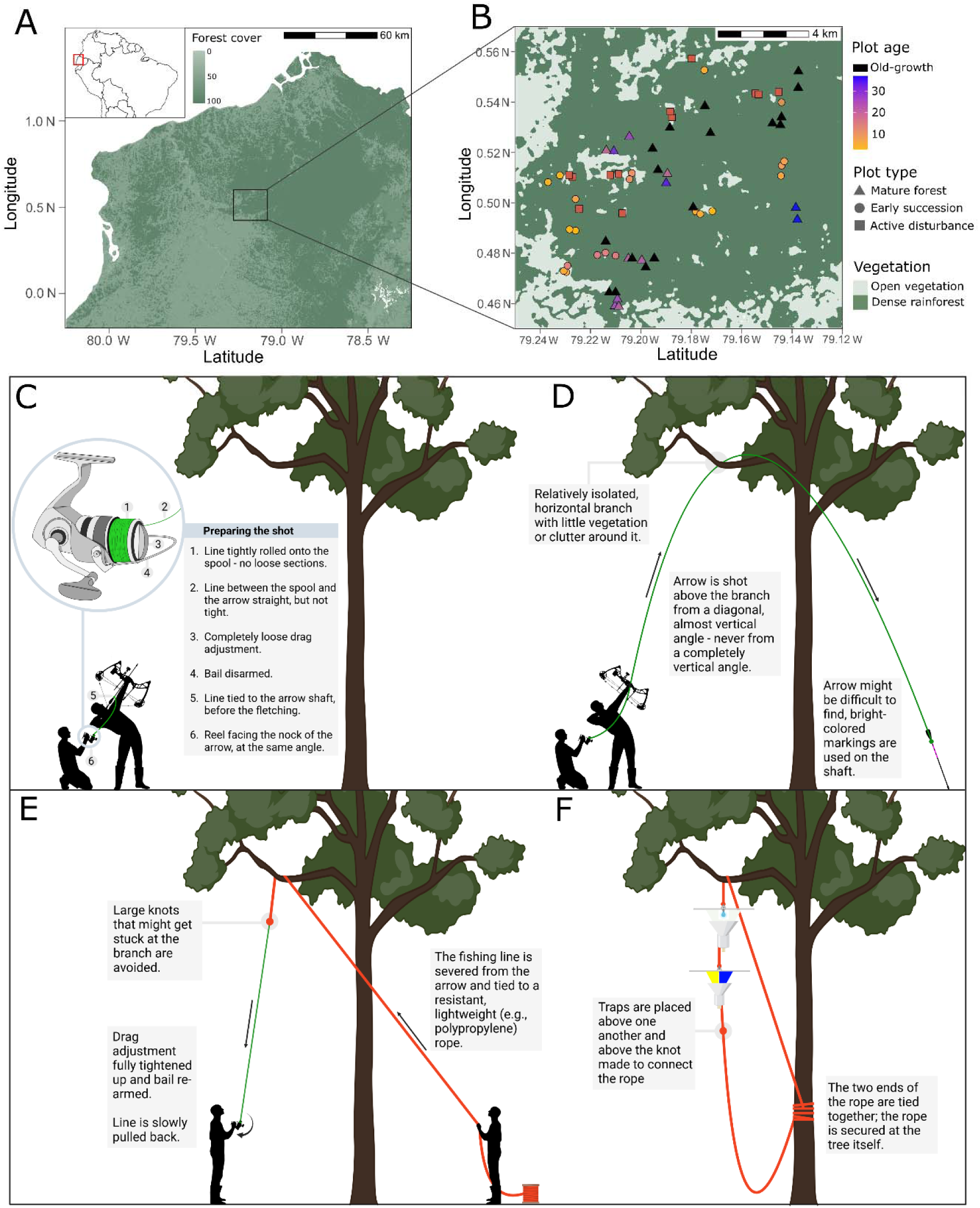
A-B: Research sites and sampling plots. C-D: A step-by-step process of installing the canopy rope that constitutes the pulley system used to lift the traps. A: location of the research area in northwestern Ecuador. B: Plots where pollinator and interaction data were collected. Plots are colored according to their recovery ages, and different shares represent recovery stages. C: preparations for the shot and setup of the fishing reel and arrow. The inset box outlines the technical requirements for a successful shot. D: Selection of a fitting branch at the canopy, shooting, and arrow retrieval. See Box S1 for a list of potential issues and their respective solutions in this step. E: Replacement of the fishing line with the final pulley rope. F: Overview of the successfully set rope and traps. Persons, traps, and trees are not to scale. Created with elements from BioRender (https://BioRender.com/i24e291).

The plots are divided into 12 active disturbances (active cacao plantations or pastures), 19 early recovery plots marked by the absence of canopy cover (1-14 years of recovery), 14 late recovery plots (15-38 years), and 17 old-growth forests with no evidence of recent disturbance. Plots were separated by a minimum distance of 350 m and a maximum distance of 12.8 km (Fig. 1B). Four insect sampling campaigns were carried out in total, with each plot sampled twice: two campaigns in 2022, and two campaigns in 2023 (half of the plots were sampled between March and May and the respective other half between October and December of each year). All plots were sampled in the understory, and late recovery and old-growth plots were also sampled in the canopy. Five late recovery plots (ranging from 15 to 26 years) could not be sampled in the canopy due to unfavorable conditions for setting canopy pulley ropes (see below). The permit to conduct research on the site and to sample insects was granted by the Ministry of Environment of Ecuador (Contrato Marco MAE-DNB-CM-2021-0187).

### Canopy pulley system

In order to sample the canopy of late-successional and old-growth forests, we installed a pulley system in a single canopy tree with the assistance of bow-and-arrow shooting (Fig. 1C-F). For the rope installation, we selected an accessible and somewhat horizontal branch from the tallest or among the tallest trees within the plot (mean height 25.8 ± 5.3 m, median 25.2 m, range 15–37.0 m) (Fig. S1). We used a 29-inch glass fiber and aluminum Strongbow GECKO compound bow (BogenSportWelt, Germany) and 70 cm glass fiber arrows with rounded points (Toparchery, Germany). A 0.32 mm 4-strands braided polyethylene fishing line (Mounchain, Japan) was attached to the arrows and set onto a TP500 double-drag brake system fishing reel (YJIU, China). Once successfully placed, the fishing line was replaced by a 5 mm polypropylene braided plastic rope, the final pulley rope to which traps were attached. A troubleshooting guide for the canopy setting is available in the Supplementary Information (Box S1).

### Pollinator sampling

We employed three different trap types in all 62 understories and all 26 canopies to capture a variety of pollinator groups. In 2022, we used nocturnal white vane traps equipped with mixed white and UV LED lights [^28^] to capture moths and nocturnal bees (Fig. S1A, Box S2 for trap details). All nocturnal bees (all belonging to the Megalopta genus, Halictidae) were taken and processed, while for moths, two focal groups were chosen for further processing and analysis: Hawkmoths (Sphingidae) and tiger moths (Arctiinae). The former is a frequently studied and key pollinator group with a functional morphology highly adapted to floral visitation [^29^], while the latter is a highly diverse group of often aposematic moths that have recently been suggested to play a key role in pollination [^30^]. Alongside nocturnal traps, we also set diurnal blue-yellow vane traps [^31^] to attract diurnal bees (Fig. S1B, Box S2). All bees were kept for processing. Both traps were set once in each plot and stratum and were left for 24h. Unlike regular vane traps, both models were adapted with a dry-killing chamber that allowed for the collection of pollen samples.

In 2023, diurnal vane traps were set again in all plots to increase bee capture numbers. In addition, fragrance traps were set to capture male orchid bees, a prominent group of pollinators in the Neotropics [^32^] (Fig. S1C, Box S2). Four fragrance traps were set at a time, each containing a different compound to capture species with different affinities towards fragrances: 1,8-cineole, eugenol, methyl salicylate, and skatol. In total, 616 traps were set (88 light vane traps, 176 colored vane traps, and 352 fragrance traps), of which 182 were set in the 26 canopies of late recoveries and old growths.

### Insect determination and trait measurements

After sorting, insects were identified to the lowest possible level. Moths were identified using the collections in the Phyletisches Museum Jena, Germany and the research collection of GB from Costa Rica, Ecuador, and Perú, as well as photographs taken by GB in the Natural History Museum (London). Identification supported by COI barcoding of ca. 90% of all moth species [^33^]. Nearly all moth species could be assigned a genus, and most were identified to species level. Bees were identified using specialized literature [^32,34,35^]. All bees could be assigned a genus and most (>90%) a species.

From each species, we collected three morphological traits: body size (BS), relative proboscis length (RPL), and wing aspect ratio (AR). BS is an indicator of dispersal ability and resistance to desiccation in insects [^36^], and was measured as the length between the tip of the head and the tip of the last abdominal segment of moths. The intertegular distance was used as a proxy for size in bees [^37^]. Proboscis length is related to morphological coupling with flowering plants and is an interaction trait associated with feeding specialization [^38^]. RPL It was measured as the proboscis length (combined extended length of the prementum and glossa in bees, and the entire extended length of the galeae in moths) divided by BS to account for the allometric relationship between proboscis length and body size in insects [^39^]. AR is a measure of flight performance, with a higher AR indicating longer and narrower wings, which serve as a proxy of gliding capability and flight speed, and may influence habitat use [^40^]. It was measured as the ratio between the square root of the forewing length and the wing area for moths; for bees, the formula was simplified to the ratio between wing length and the wing’s mean chord (widest wing width) due to a relatively conserved wing shape across groups [^40^]. Traits were measured manually with a caliper for larger bees (e.g., Euglossini) and with the aid of microscopy and linear morphometric analysis via the ImageJ software [^41^] for smaller (> 5 mm of body length) bees. For the latter, the proboscises were dissected and mounted on slides for further image analysis. All moth traits were collected via standardized photography and processed using image analysis, with a modified version of the mothseg algorithm [^42^]. We supplemented this data with individual measurements of photographs with scale bars for proboscis length. Up to ten individuals were measured per species.

In moths, we measured two additional and important traits related to coloration, which is linked to thermoregulation and predation avoidance [^43^] and thus potentially a driver of the vertical distribution of species. For both, we also used standardized moth photographs and the modified version of the mothseg algorithm: (i) Lightness was measured as the median pixel value of four main color channels (red, green, blue, and UV) of the dorsal side of fully spread individuals, which represents overall brightness (e.g., high values for white species and low values for dark species), (ii) Contrast, measured as the interquartile distance (25th-75th percentiles) of the distribution of pixel values, representing coloration variability with high values often associated with conspicuousness to predators and aposematism.

### Pollen DNA metabarcoding

After capture, all insects were checked under a stereomicroscope for the presence of pollen, and any significant pollen load (10 or more grains) was collected. In bees, pollen was collected both from the specialized structures of females (entire legs from corbiculate species or by scraping the pollen out of scopae), or, when present, by scraping other body surfaces, such as the head, thorax, and abdomen, of both males and females. In male orchid bees, the entire pollen sacs deposited by orchids were collected. As for moths, due to the large number of individuals in the traps and a higher chance of contamination, only proboscises, which remain coiled between the palps, were checked for pollen and collected. Samples were pooled per pollinator species and per plot, and preserved dry at −20°C.

Sample extraction, amplification, and sequencing were performed at Ludwig-Maximilians-Universität München, Germany. Extraction, quality control, and library preparation were done as described in [^44^] and [^45^], followed by multiplexed next-generation DNA metabarcoding for the ITS2 region [^46^]. The bioinformatics pipeline is available at [^47^] (version 8c8536b). We used VSEARCH [^48^] for quality filtering, merging, dereplication, and definition of amplicon sequence variants (ASVs). We first classified ASVs through global alignments (97% threshold) against an Ecuadorian plant database using BCdatabaser [^49^]. We then identified unclassified ASVs using a global vascular plant database [^50^] with VSEARCH, again at a 97% threshold. We hierarchically identified the remaining unclassified reads to the lowest possible taxonomic level at a threshold of 0.8 with SINTAX [^51^]. Contaminations (e.g., Eurasian or cultivated genera not found in the area) and other potential contaminations observed in the negative controls were removed with phyloseq [^52^]. Finally, we merged ASVs at the species level and transformed them into relative read abundances (RRAs) per sample. Following [^30^], the throughput of each pool was multiplied by the number of individuals in the pool (median = 1, range = 1–65), then randomly subsampled and unevenly rarefied based on sample size, before being multiplied again to account for uneven pool sizes that may affect sequence detection probabilities. Low RRAs (<10%) were then removed from the samples according to the results of the positive control samples to further exclude possible contamination or non-legitimate interactions (e.g., low-abundance pollen deposited on pollinators by wind). RRAs were then used as interaction weights between plant species and pollinator species per sample, which avoids overemphasizing rare species in pollen pools [^30^]. Final interaction weights thus corresponded to the rarefied RRAs multiplied by 1000 to obtain integers that approximate observation counts.

### Data analysis

#### 1. Response variables

All data were analyzed in R 4.5.0 [^53^], using the packages vegan for community-level analyses [^54^], bipartite for network analyses [^55^], FD for estimating functional richness [^56^], and betapart for estimating beta-diversity [^57^]. The following response variables from pollinator communities were measured per sample (each pollinator group was analyzed independently):

i. Abundance: the total sum of individuals.
ii. Alpha-diversity: measured as the exponential Shannon diversity (eH’) (effective diversity, q = 1), which corresponds to the number of equally common species that would yield the same diversity value [^58^].
iii. Beta-diversity: measured by two parameters, β_DISTINCTIVENESS_ and β_DISPERSAL_. Both metrics are based on a non-metric multidimensional scaling (NMDS) (k = 2, minimum of 10000 iterations) of the samples based on a Bray-Curtis distance matrix. β_DISTINCTIVENESS_ was measured as a sample’s mean total abundance-weighted dissimilarity in relation to all other samples, being a proxy for the uniqueness of the sample’s composition in relation to the entire sample pool [^57^]. β_DISPERSAL_ was measured as the distance of each sample to its within-group centroid (i.e., recovery status and stratum), indicating whether a certain sample group has a conserved and predictable species composition or shows high variation [^57^].
iv. Functional richness: measured via a distance-based and multi-trait framework. Functional richness represents the volume of the minimum convex hull encompassing all species in a Principal Correspondence Analysis reduction based on community-weighted trait means [^59^]. High functional richness thus represents a sample where species occupy a large area of the trait space. All bee traits (RPL, BL, AR) and all moth traits measured (RPL, BS, AR, lightness, and coloration) were employed. The index could only be calculated for samples with three or more species.
v. Community-weighted traits means (CwMean) and coefficient of variation (CwCV): in order to also examine trait-specific responses, we measured mean values and coefficient of variation of pollinator traits (RPL, BS, AR, lightness, and contrast) in each sample, weighted by species abundances. Additionally, the following variables were measured for each plot’s interaction network (here, interaction networks were built with all pollinator groups combined):
vi. Interaction diversity (H’): Shannon’s diversity of network entries.
vii. Interaction specialization: measured as the H_2_’ index of niche differentiation [^60^]. H_2_’ varies from zero to one and quantifies how unique, on average, the interactions between species are. High values (H_2_’ → 1) indicate a low interaction overlap between pollinator species and thus high network specialization.
viii. Interaction beta-diversity: measured as in (iii). The dissimilarity matrix was constructed based on an occurrence matrix, where species corresponded to unique interaction pairs found in each sample, and their weights represented their abundance. Dimensionality reduction and β_DISTINCTIVENESS_ and β_DISPERSAL_ calculation were then performed accordingly.

#### 2. Independent variables and linear models

Plot recovery age (years after the last disturbance) and stratum (understory or canopy) were used as explanatory variables for the variables above along the recovery gradient. Because old-growth forests do not have a determined age, we transformed plot recovery age into the discrete variable recovery status by separating plots into the following groups: active disturbance (0, plots still undergoing disturbance), early regeneration (1, plots between one and 14 years of regeneration), late regeneration (2, plots between 15 and 38 years of regeneration), and old growth (3). Besides these two main variables, we also included potential landscape features that might affect insect abundance and diversity patterns: plot elevation a.s.l., which may affect plot temperature and thus the microclimatic niches in both strata, and the proportion of forest within a 500 m buffer around the plot, a proxy for landscape isolation and fragmentation, which may affect species pools due to dispersal limitations. The ESRI World Imagery basemap was used as a source for recent landcover and sourced from Escobar et al. (2024).

Data were analyzed in two ways: first, we examined the paired differences in pollinator community parameters, interaction network parameters, and community-weighted traits in canopy-bearing forests (late successional and old-growth) to first determine whether parameters differ between forest strata and/or recovery legacies. Linear mixed-effects models (LMMs) were fitted for each response variable, setting stratum (canopy vs. understory) and recovery stage (successional vs. old growth) as independent variables and collection plot as a random variable to account for paired samples. Models were fit as generalized mixed-effect models (GLMMs) with a Poisson error family in all instances of pollinator abundance, and with a Gaussian error fit (regular LMMs) in all other instances. Residual normality and homoscedasticity were checked for all models. Second, all chronosequence samples (62 understory and 26 canopy samples) were analyzed together to assess the effect of stratification, recovery status, and their interaction on pollinator community and interaction network reassembly. Similarly, LMMs were fitted for each variable, with stratum, recovery status, and their interaction as independent variables, alongside plot elevation and the proportion of surrounding forest. Again, the collection plot was set as a random variable. Error family selection, as well as residual normality and homoscedasticity checks, were performed as above.

## RESULTS

### 1. Pollinators sampled

A total of 20,245 target insects from 424 species were collected in the 88 samples (62 understories and 26 canopies): 8,615 bees of 86 species and 11,630 moths of 338 species. Moths were represented by the two focal groups Arctiinae and Sphingidae, with the former accounting for the vast majority of individuals (98.1%). Therefore, all results concerning moths hereafter refer to both groups together. The samples from diurnal vane traps consisted almost exclusively of stingless bees (Apidae: Meliponini) (3,607 individuals of 29 species), while orchid bees (Apidae: Euglossini) were the only group collected by fragrance traps (4,530 individuals of 53 species). Nocturnal Megalopta (Halictidae: Augochlorini) bees were collected from light traps (488 individuals of four species). Other bee groups (e.g., Colletidae, diurnal Halictidae) accounted for 0.27% of the total bee captures and were thus not included in further analyses.

### 2. The role of stratification and disturbance legacy in mature forests

In paired canopy and understory samples from mature forests (old growth and late successional forests), stratification was a stronger driver of pollinator communities than recovery legacy (Tab. 1, Fig. 2). Abundance, alpha diversity, functional richness, and beta diversities varied significantly between canopy and understory in all groups, but rarely between old-growth and successional forests (Tab. 1). Moths and nocturnal bees were more abundant and diverse in canopies (Fig. 2A, D, Tab. 1). Nocturnal bees were additionally more abundant in old-growth forests (Fig. 2D). Moth beta diversity, however, was not affected by none of the two predictors (Fig. 3C, Tab. 1).

**Figure 2.**
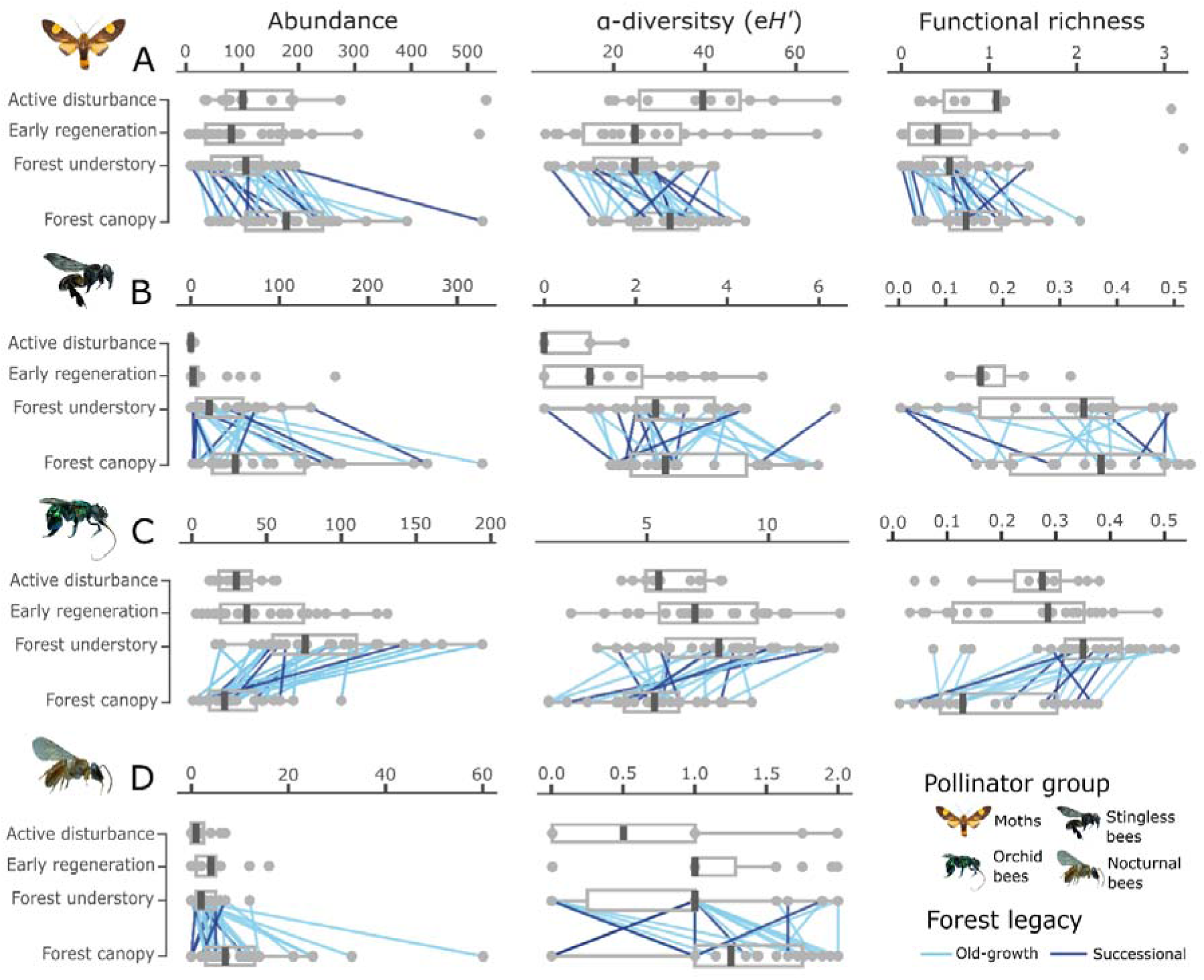
The abundance (left), alpha diversity (exponential Shannon diversity) (middle), and functional richness (right) of different diurnal and nocturnal pollinator groups along a forest recovery chronosequence and between forest strata. A: tiger moths, B: stingless bees (Meliponini), C: orchid bees (Euglossini) and D: nocturnal bees (Megalopta genus). Forests (plots older than 15 years and with a clear vertical structure) are separated into understory and canopy samples. Lines between points indicated paired samples from the same plot, and line color represents disturbance legacy (old growth in light blue and successional forests between 15 and 38 years of age in dark blue). Functional richness and values are not shown for samples with fewer than three species, as was always the case with nocturnal bees.

**Figure 3.**
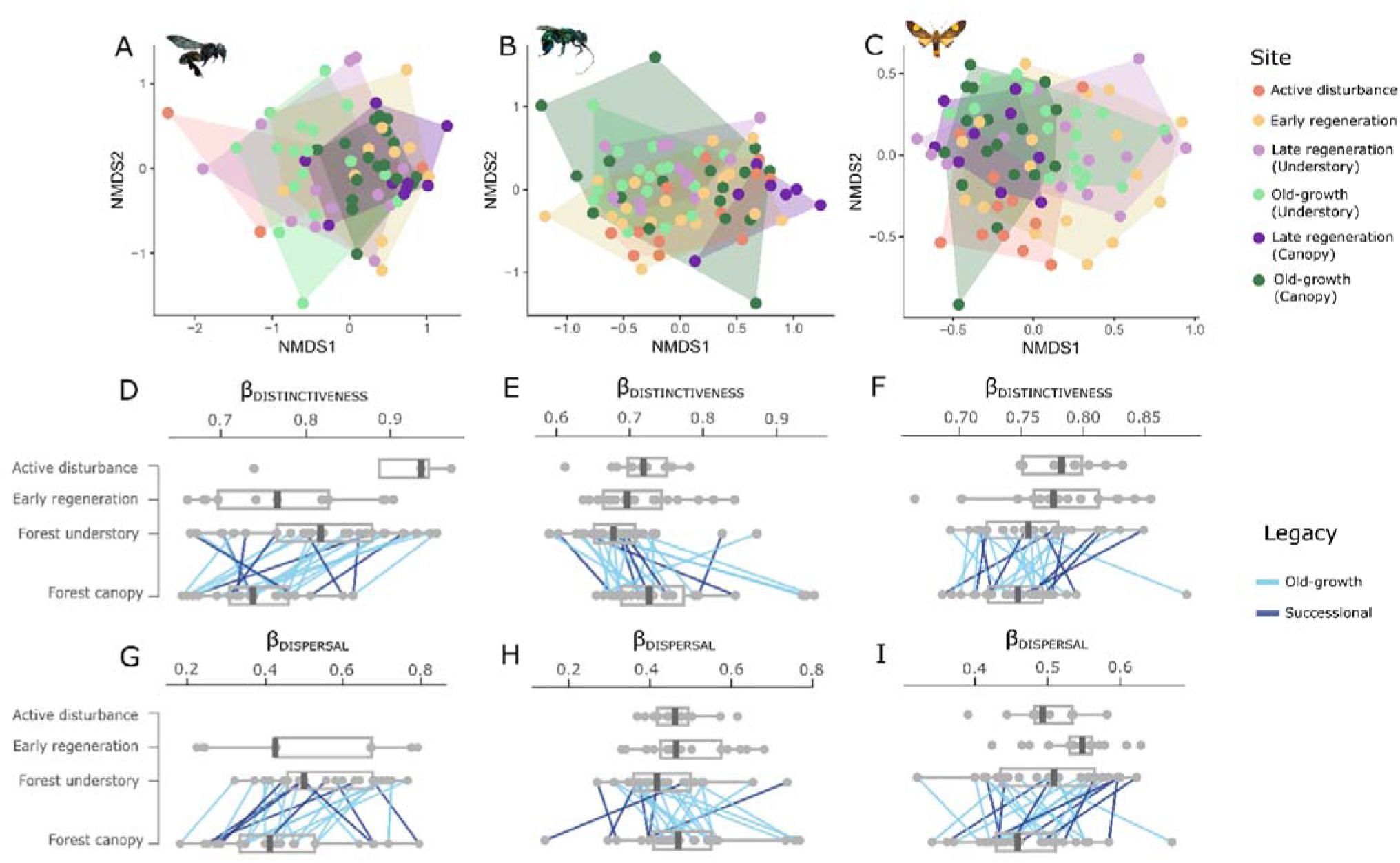
The composition of pollinator communities in understory and canopy samples across the recovery chronosequence. On the top row (A-C), the ordination of samples according to pollinator communities based on non-metric multidimensional scaling (NMDS). The first two axes are shown. Samples are colored according to recovery status and, in late successional forests, are divided into canopy and understory. A – stingless bees (Meliponini), B – orchid bees (Euglossini), C – moths. On the middle row (D-E), values of beta diversity are shown per sample in terms of distinctiveness (mean distance to shared centroid). On the bottom row (G-I), values of beta diversity are shown per sample in terms of dispersal (within-group mean distance from centroid). In D-I, samples are divided by recovery status, and forest samples (old growth and successional) are paired between canopy and understory.

**Table 1.** The role of stratification and disturbance legacy as drivers of pollinator abundance, alpha diversity (Rarefied eH’), and functional richness (R) in paired undergrowth and canopy samples in vertically structured tropical forests (>15 years). The mixed-effect linear models’ marginal R-squared is shown (_m_R^2^), as well as the SD-standardized estimate coefficients (β) and standard errors (SE) for each explanatory variable, i.e., (canopy vs. understory) and disturbance legacy (old growth vs. late successional). The fit used for each model’s error family is specified. Bold values denote significance. *p < 0.05, **p < 0.05, ***p < 0.001.

| | Error fit | $mR^2$ | Canopy vs. Understory | Old growth vs. Successional |
| --- | --- | --- | --- | --- |
| | | | $\beta$ (SE) | $\beta$(SE) |
| **Moths** |  |  |  |  |
| Abundance | Poisson | 0.24 | **0.003 (0.0001)** *** | 0.0002 (0.001) |
| α-diversity ($eH'$) | Gaussian | 0.20 | **0.39 (0.10)** *** | -0.05 (0.16) |
| Functional R | Gaussian | 0.13 | **0.37 (0.11)** ** | 0.02 (0.16) |
| β-Dispersal | Gaussian | 0.05 | -0.20 (0.13) | -0.12 (0.15) |
| β-Distinctiveness | Gaussian | 0.03 | -0.11 (0.13) | -0.14 (0.15) |
| **Stingless bees** |  |  |  |  |
| Abundance | Poisson | 0.23 | **0.006 (0.0003)** *** | 0.005 (0.003) |
| α-diversity ($eH'$) | Gaussian | 0.06 | 0.12 (0.12) | 0.21 (0.16) |
| Functional R | Gaussian | 0.03 | 0.18 (0.12) | 0.002 (0.19) |
| β-Dispersal | Gaussian | 0.10 | **-0.33 (0.10)** *** | 0.008 (0.17) |
| β-Distinctiveness | Gaussian | 0.19 | **-0.44 (0.12)** *** | 0.03 (0.14) |
| **Orchid bees** |  |  |  |  |
| Abundance | Poisson | 0.59 | **-0.01 (0.001)** *** | 0.001 (0.002) |
| α-diversity ($eH'$) | Gaussian | 0.23 | **-0.49 (0.13)** *** | 0.02 (0.13) |
| Functional R | Gaussian | 0.34 | **-0.56 (0.11)** *** | -0.18 (0.14) |
| β-Dispersal | Gaussian | 0.15 | 0.19 (0.14) | **0.35 (0.14)** * |
| β-Distinctiveness | Gaussian | 0.14 | **0.39 (0.14)** ** | -0.01 (0.14) |
| **Nocturnal bees** |  |  |  |  |
| Abundance | Poisson | 0.50 | **0.06 (0.006)** *** | **0.03 (0.01)** * |
| α-diversity ($eH'$) | Gaussian | 0.13 | **0.30 (0.12)** * | 0.22 (0.14) |

Stingless bees were more abundant in canopies, but not more diverse or functionally rich (Tab. 1, Fig. 2 B). Moreover, their communities were more conserved in canopies, being less distinct and exhibiting less dispersal than those in the understories (Tab. 1, Fig. 3A).

In contrast to other groups, orchid bees showed an opposite pattern, being strongly associated with the understory in abundance, diversity, and functional richness, with canopies presenting the most distinctive communities (Tab. 1, Fig. 2C, Fig. 3B). Communities of these bees in old-growth forests also presented higher dispersal (Tab. 1, Fig. 3B).

### 3. The effect of stratification in pollinator community reassembly

In the context of the chronosequence, stratification and recovery status were significant drivers of the abundance of all pollinator groups. Stingless bees were more abundant in canopies of older forests, as well as more diverse in older forests in general, while nocturnal bees were both more abundant and diverse in the canopies of older forests (Tab. 2, Fig. 2 B, C). None of these two groups was affected by the amount of surrounding forest or elevation.

**Table 2.** Drivers of abundance, alpha diversity (exponential Shannon diversity – eH’), functional richness (R), and beta diversity (dispersal and distinctiveness) of different diurnal and nocturnal groups along a tropical forest recovery chronosequence. Drivers: Stratification (Understory vs. Canopy); Recovery: recovery status (active disturbance, early regeneration, late regeneration forest, or old-growth forest); Rec:Strat: the interaction between stratification and recovery status; Elevation: elevation a.s.l.; ForestArea: plot isolation (proportion of forest within a 500m buffer). The mixed-effect linear models’ marginal R-squared is shown (_m_R^2^), as well as the SD-standardized estimate coefficients (β) and standard errors (SE) for every explanatory variable in each model. Bold values denote significance. *p < 0.05, **p < 0.01, ***p < 0.001.

| | Error fit | $mR^2$ | Stratification<br>$\beta$ (SE) | Recovery<br>$\beta$ (SE) | Strat:Rec<br>$\beta$ (SE) | Elevation<br>$\beta$ (SE) | ForestArea<br>$\beta$ (SE) |
| --- | --- | --- | --- | --- | --- | --- | --- |
| **Moths** |  |  |  |  |  |  |  |
| Abundance | Poisson | 0.30 | **0.006 (6x10<sup>-4</sup>)\*\*\*** | **-0.003 (0.001)\*\*** | **-0.004 (6x10<sup>-4</sup>)\*\*\*** | -0.001 (0.001) | **0.004 (0.001)\*\*\*** |
| α-diversity ($eH'$) | Gaussian | 0.21 | 0.03 (0.41) | **-0.61 (0.15)\*\*\*** | 0.26 (0.41) | 0.10 (0.12) | **0.39 (0.15)\*** |
| Functional R | Gaussian | 0.19 | -0.14 (0.43) | **-0.46 (0.15)\*\*** | 0.40 (0.43) | 0.23 (0.12) | **0.33 (0.15)\*** |
| β-Dispersal | Gaussian | 0.20 | -0.92 (0.49) | 0.05 (0.14) | 0.72 (0.50) | **0.31 (0.11)\*\*\*** | **-0.34 (0.13)\*\*** |
| β-Distinctiveness | Gaussian | 0.20 | -0.76 (0.49) | -0.07 (0.14) | 0.70 (0.50) | 0.11 (0.11) | **-0.38 (0.14)\*\*** |
| **Stingless bees** |  |  |  |  |  |  |  |
| Abundance | Poisson | 0.62 | 0.001 (0.002) | **0.02 (0.004)\*\*\*** | **0.005 (0.001)\*\*** | 0.005 (0.003) | 0.004 (0.004) |
| α-diversity ($eH'$) | Gaussian | 0.38 | -0.19 (0.41) | **0.46 (0.12)\*\*\*** | 0.30 (0.41) | -0.11 (0.10) | 0.15 (0.12) |
| Functional R | Gaussian | 0.15 | 0.43 (0.67) | 0.23 (0.18) | -0.25 (0.69) | 0.11 (0.17) | 0.11 (0.18) |
| β-Dispersal | Gaussian | 0.06 | 0.06 (0.43) | -0.12 (0.34) | 0.20 (0.42) | 0.09 (0.16) | 0.11 (0.18) |
| β-Distinctiveness | Gaussian | 0.18 | 0.48 (0.50) | -0.09 (0.15) | -0.89 (0.51) | -0.09 (0.13) | 0.22 (0.16) |
| **Orchid bees** |  |  |  |  |  |  |  |
| Abundance | Poisson | 0.40 | **-0.01 (0.002)\*\*\*** | **0.008 (0.002)\*\*\*** | -0.002 (0.003) | -0.004 (0.002) | 0.004 (0.003) |
| α-diversity ($eH'$) | Gaussian | 0.17 | -0.38 (0.56) | 0.32 (0.24) | -0.11 (0.57) | 0.02 (0.11) | -0.10 (0.57) |
| Functional R | Gaussian | 0.24 | 0.51 (0.51) | 0.16 (0.14) | -1.04 (0.52) | -0.04 (0.11) | **0.28 (0.14)\*** |
| β-Dispersal | Gaussian | 0.21 | **-1.86 (0.53)\*\*\*** | -0.01 (0.13) | **2.14 (0.54)\*\*\*** | 0.15 (0.11) | **-0.30 (0.13)\*** |
| β-Distinctiveness | Gaussian | 0.19 | 0.08 (0.54) | -0.07 (0.14) | 0.36 (0.55) | 0.20 (0.11) | -0.23 (0.13) |
| **Nocturnal bees** |  |  |  |  |  |  |  |
| Abundance | Poisson | 0.40 | **-0.17 (0.04)\*\*\*** | 0.006 (0.02) | **0.23 (0.04)\*\*\*** | 0.024 (0.01) | -0.002 (0.02) |
| α-diversity ($eH'$) | Gaussian | 0.16 | -0.75 (0.48) | 0.12 (0.15) | **1.02 (0.49)\*** | -0.14 (0.12) | -0.03 (0.14) |

Orchid bees, conversely, were significantly less abundant in canopies in general but associated with late successional forests and old growth (Tab. 2, Fig. 2C), and the canopies of older forests had the most distinct communities (Tab. 2, Fig. 3B). The amount of surrounding forest also affected these bees, with community composition presenting less variation and higher functional richness in plots near higher proportions of forest (Tab. 2).

Moths, despite being associated with canopies, were also abundant and diverse in active disturbances and early regenerating forests, therefore being significantly associated with early successional understories when inserted into a chronosequence context (Tab. 2, Fig. 2). The surrounding forested area was a strong driver of all moth community parameters, with moths being more abundant, diverse, and functionally rich in more connected plots, while presenting more conserved and predictable communities (Tab. 2, Fig 3C).

### 4. The role of functional traits

Forest recovery and stratification had different effects on the functional traits of pollinators. Moths were the least affected functionally by both variables, as most traits did not vary between recovery stages or between canopy and understory (Fig. 4A). However, older canopies contained moths with a significantly higher aspect ratio (narrow and long wings), and canopies in general presented moths with more variable contrasts in coloration.

**Figure 4.**
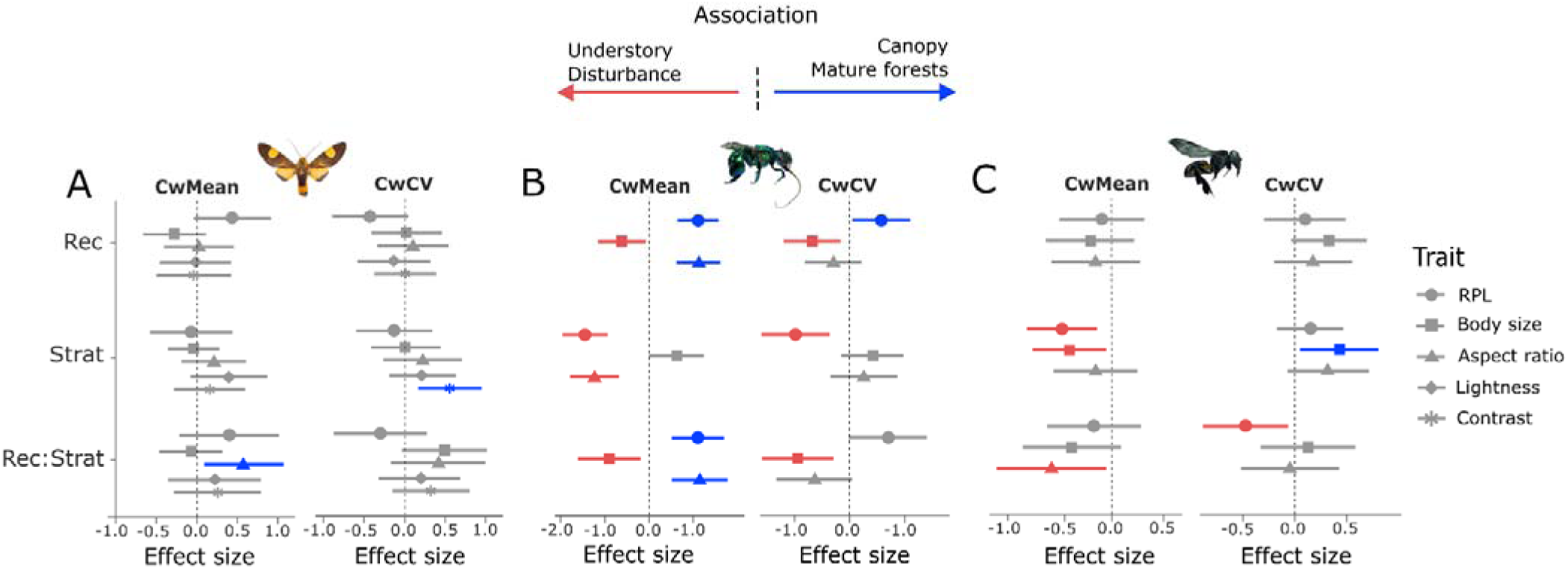
Effect sizes (standardized estimate coefficients) of recovery stage (Rec), stratification (Strat), and their interaction (Rec:Strat) on the community-weighted means (CwMean) and community-weighted coefficient of variation (CwCV) of different functional traits from pollinators. A: moths, B: orchid bees (Euglossini), C: stingless bees (Meliponini). The mean effect size and 95% confidence intervals are shown. Significant effects are shown in blue (positive association with canopies and older plots) or red (positive association with understories and disturbance or early regeneration).

Orchid bees were the group whose functional morphology was most affected by both variables (Fig. 4 B). Bees in older canopies had relatively longer proboscises, had higher aspect ratios, and were smaller. Additionally, bees had more variable proboscis lengths in older forests and in understories. Body size also varied more frequently in the understories of younger forests or disturbances.

Finally, the functional traits of stingless bees did not vary in function of the recovery stage, but rather stratification or its interaction with recovery stage (Fig. 4 C). Bees were larger and had relatively shorter proboscises in the understory, but body size was more variable in canopies. Moreover, aspect ratios were larger in the understories of younger forests, as well as the variation in proboscis length.

### 5. Reassembly of interaction networks

Stratification had significant effects on plant-pollinator interactions. Paired samples from canopy-bearing forests showed that interaction diversity was higher in canopies (R^2^ = 0.11, β = 0.3, se = 0.1, p < 0.01) (Fig. 5A) but not network specialization (R^2^ = 0.01, β = −0.02, se = 0.1, p = 0.08) (Fig. 5B). In terms of beta-diversity, interactions were the most distinct in canopies (R^2^ = 0.49, β = 0.67, se = 0.1, p < 0.001) (Fig. 5C) but not more dispersed (R^2^ = 0.05, β = −0.19, se = 0.13, p = 0.15) (Fig. 5D). Forest legacy did not affect any of the parameters in paired forest samples (p > 0.05 in all instances).

**Figure 5.**
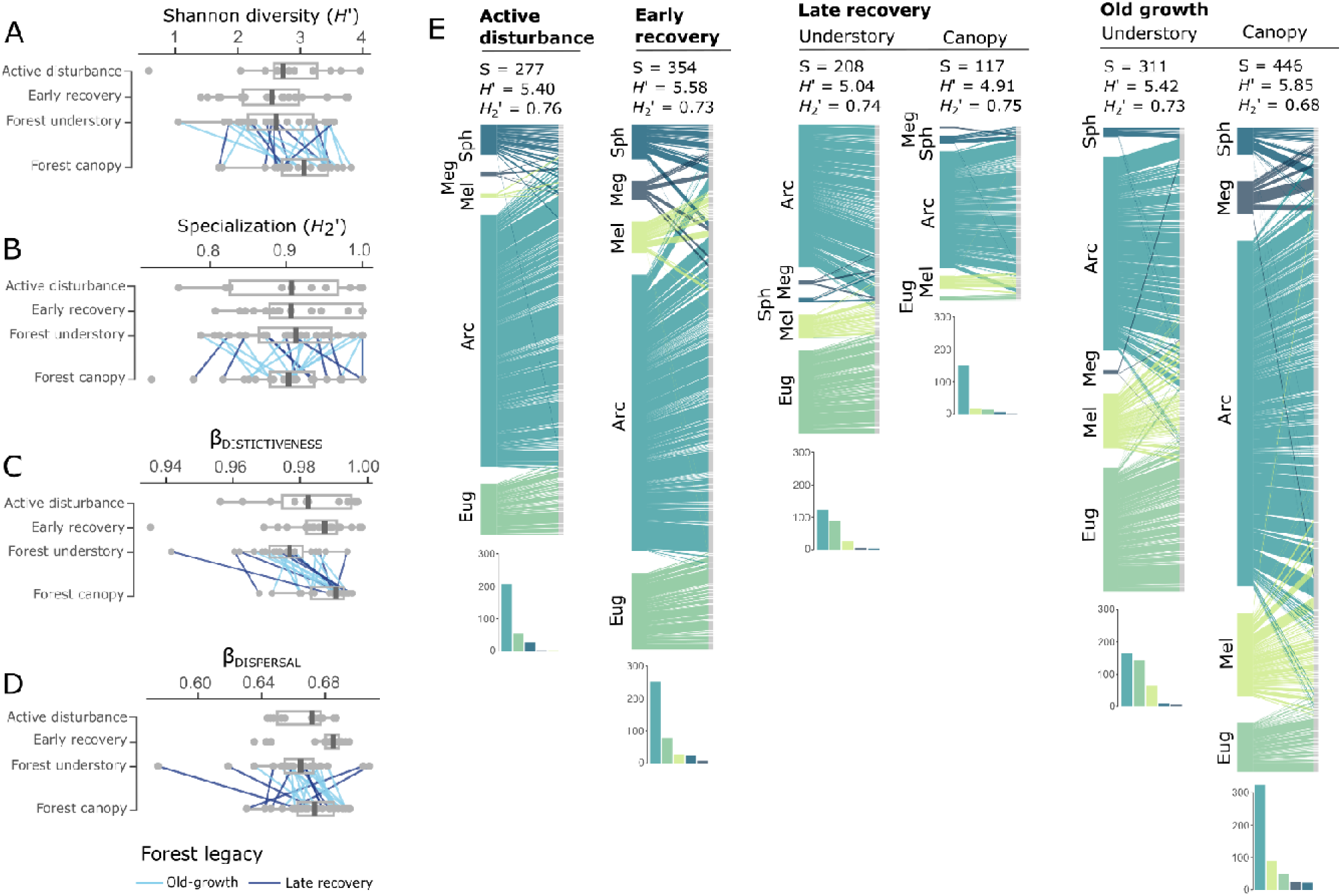
The diversity and structure of Interaction networks between pollinators and plants along the recovery gradient and forest strata. A – plot-level interaction Shannon diversity (H’), B – Network specialization (H_2_’), C – interaction distinctiveness (beta diversity measured as mean distance to shared centroid), and D – Interaction dispersal (beta diversity measured as within-group mean distance from centroid). E – The structure of networks formed by interactions pooled by the recovery stage and stratum, followed by network size (S), interaction Shannon diversity (H’) and network specialization (H_2_’). Pollinators and their interactions are colored according to major groups (Sph: Sphingidae, Meg: Megalopta, Arc: Arctiinae moths, Mel: Meliponini bees, Eug: Euglossini bees). At the bottom of each network, bar plots show the frequency of pollen samples collected for each pollinator group. Although networks in E are illustrated with pooled pollinator groups, all metrics in A-E were measured in species-level networks (pollinator species and plant ASVs).

When setting interactions within the chronosequence setting, networks were primarily driven by landscape features. Interaction diversity was positively affected by the proportion of surrounding forest only (R^2^ = 0.18, β = 0.42, se = 0.15, p < 0.01). Network specialization (R^2^ = 0.13) was higher in higher elevations (β = 0.26, se = 0.13, p < 0.05) and in sites with less surrounding forest (β = −0.40, se = 0.15, p < 0.05). As for beta-diversity, interactions were the most distinct in plots with less surrounding forest (R^2^ = 0.19, β = −0.38, se = 0.14, p < 0.005), and more dispersed (R^2^ = 0.20) at higher elevations (β = 0.31, se = 0.11, p < 0.005) and in plots with less surrounding forest (β = −0.34, se = 0.13, p < 0.05). Stratification and recovery status, as well as their interaction, did not have significant effects on any of the parameters within the chronosequence (p > 0.05 for all estimates).

Pooled networks were stable across recovery statuses and strata, remaining similar in terms of diversity and specialization despite clear changes in size (Fig. 5E). Late-recovery networks were the smallest and dominated by settling moths, particularly the canopy. Old-growth forests contained the largest, least specialized networks, as well as the highest diversity of pollinator groups. The old-growth canopy network was marked by its large size and the participation of meliponine bees, nocturnal bees, and hawkmoths while maintaining the dominance of settling moths (Fig. 5E).

## DISCUSSION

The vertical stratification of resources in forests is a key driver of the distribution of various animal functional groups, ranging from non-flying mammals [^61^] to butterflies [^62^], as well as ground predators and flying insectivores [^63,64^]. Resource stratification thus permeates different ecosystem processes. For pollinators, floral resource stratification has been reported to influence the abundance and diversity of bees and wasps in temperate forests [^65,66^] and of moths in Paleotropical forests [^67^], for instance. Moreover, large, highly energy-dense canopy trees sustain populations of tropical bees and beetles [^9,20,26,62^], and an overall higher diversity of flowering resources has been recorded in tropical canopies compared to understories [^7–10^]. These premises suggest that canopies are essential to sustain pollinator populations in the landscape, especially in highly disturbed areas with decreasing forest cover. However, the role of canopies in the reassembly of pollinator communities and their interaction networks along forest recovery and succession had thus far not been explored.

Our study revealed that all pollinator groups studied exhibited some degree of stratification in their abundance, diversity, and functional richness within the chronosequence. Most groups showed an association with canopies, except for the understory-associated orchid bees. In canopy-bearing forests (successional and old-growth), forest stratum was a stronger predictor of pollinator community parameters than forest legacy, suggesting that resource stratification towards the canopy (e.g., flowers, host plants, or nesting sites) may already occur within a few recovery decades. The canopy-closure phase of succession (10-25 years) is characterized by rapid species turnover, an increase in basal area, maturity of pioneer trees, exclusion of understory pioneers, and the establishment of an epiphytic flora [^69^], all of which may influence pollinator stratification. Even though several veteran tree species may take centuries to reestablish, leading to distinct communities in old-growth forests compared to those in successional [^12^], sufficient resource density may already be present in old secondary forests to sustain pollinators [^70^], and these resources seem to stratify rather soon.

However, contrary to trends observed in pollinator communities, interaction networks were large and complex in active disturbances and early-recovery plots and relatively small in successional forests. Only when both strata were combined were the networks in successional forests comparable to those of earlier stages. Old-growth forests, on the other hand, presented large and rich networks in each stratum, which led to their complete network being twice as large as those of other recovery statuses. Therefore, we can assume that pollinators are flexible in terms of foraging and diet as they are able to exploit disturbed sites within a landscape. They are likely to profit from increased herb and forb density in pastures, as well as pioneer bushes and trees in early succession (e.g., the pioneer tree genus Cecropia was the most abundant in pollen pools), potentially buffering pollinator groups against disturbance. This resistance to disturbance can be best observed in moths, which stand out as the group least affected by the recovery gradient. Tiger- and hawkmoths thrived in pastures and plantations, essentially acting as indicators of disturbance. Such resistance has been previously noted and associated with the high mobility of most species, which allows movement across the matrix during milder nighttime conditions, as well as with the feeding generalism of adults and caterpillars of many species [^42,71^]. Moths might thus be important drivers of network reassembly in recovering forests [^30^].

However, pollinators in general appear to still be associated with old-growth forests to some extent, likely due to both a more diverse pool of floral resources [^72^] and a higher likelihood of finding nesting sites and refuge [^65,73^]. Tiger moths, the leading group in the pollination networks and the one that showed the strongest affinity towards the canopy, were strongly influenced by the proportion of surrounding forests, being more abundant, diverse, and functionally richer in less isolated plots. Their communities were also more predictable and less variable in plots surrounded by more forest cover. Indeed, forest cover and plot elevation, two variables linked to plot isolation, were also the best predictors of network structure and diversity. Old-growth forests may thus provide suitable habitats and act as a source of pollinator species that are able to forage within the matrix (i.e., areas without canopy cover), which underscores the importance of maintaining forest connectivity and a minimum proportion of forest on the landscape to sustain pollinator populations [^72^]. This importance also extends to secondary forests, given the abundance and diversity of pollinators in their canopies. Despite small networks in late-recovery sites, canopy interactions were still overall more distinct and variable, suggesting that canopies in both legacies contain plant and pollinator species not observed in forest understories, indicating the presence of canopy-specialized species [see ^74^]. These species appear to be the most representative among tiger moths, which account for the majority of interactions in late-recovery canopies.

This network recovery pattern warrants a careful examination of the types of resources beyond flowers that drive the vertical distribution of floral visitors. Tiger moths, for instance, are also affected by the availability of host plants for larvae, and many species specialize in canopy-specialized lianas [^19^]. An increased (functional) diversity of host plants in canopies, driven largely by epiphytes and climbers that add to the crown cover of large trees, may thus also drive the vertical distribution and potentially the recovery of other herbivores. Canopy-associated substrates and microenvironments (e.g., nesting cavities, phytotelmas, and trunk roosts) may also influence the distribution of a plethora of other animals and their recovery patterns, encompassing small insects to vertebrates [^17,65,75,76^]. The benefits of these non-nutritional resources and microenvironmental niches are already present in late-successional forests and can enhance faunal recovery [^77^]. For pollinators specifically, the observed high resilience associated with newly formed canopies highlights the role of vertical stratification for pollinator community reassembly and further evidences the relevance of secondary forests as alternate providers of nutritional and non-nutritional resources [^70^].

Some pollinators, however, were absent in active disturbances and particularly dependent on old canopies in primary forests. The eusocial stingless bees and the facultatively social nocturnal Megalopta were positively influenced by recovery time and stratification, both of which also exhibited the most interactions in old-growth canopies. Both groups are likely limited by the availability of nesting and food resources found in forests. The former are dominant pollinators in tropical ecosystems and typically live in colonies that host hundreds to thousands of individuals [^32^]. They show a high susceptibility to forest cover loss [^78,79^], which can be attributed to the cavity-nesting habit of many species [^32^], the need for large trees (> 50 cm in diameter at breast height) to establish nests [^79^], and their reliance on forest-associated resources, such as floral or bark resin [^80^]. In our study, canopy specialists primarily consisted of small (ca. 2 mm), gregarious, and wood-nesting Trigonisca bees. These would theoretically be classified as “shade-loving” bees according to [^74^] due to their small size and resulting susceptibility to heat and desiccation. Nonetheless, stingless bees were on average smaller and had shorter proboscises in the canopy, which suggests that the interplay between functional traits and abiotic factors is not always straightforward and may interact with an animal’s need for food and nesting resources. For instance, nesting and foraging close to the canopy may enable small stingless bees with large colonies and short flight ranges (e.g., Trigonisca, less than 100 m from colonies) to compensate for lower mobility [^81^] and optimize foraging. Furthermore, establishing a nest may also be more successful in upper strata, where the abundance of natural substrate for cavity-nesting pollinators such as stingless bees and Megalopta (e.g., natural crevices in wood) is higher [^65^]. Megalopta bees also exhibit remarkably low heat tolerance [^82^], which may also be linked to their reliance on forests with canopy cover.

Although also more diverse and abundant in late-successional and old-growth forests, orchid bees stand out as relatively resistant to disturbance and remarkably canopy-avoidant, a trait previously observed before but not ubiquitous across studies [^83^]. Male orchid bees require wide dispersal and foraging ranges to search for particular and specific volatile compounds used in courtship that are sparsely distributed across the forest [^84^]. This need, combined with a good dispersal ability [^85^], may lead to a low response to the successional gradients, as bees are able to transit between forest patches. In this regard, orchid bees may contribute to the stability and reassembly of pollination networks by facilitating pollen flow across the matrix. However, the underlying causes of the repeating canopy-avoidant behavior are unknown. A prominent hypothesis is that the dissipation of floral volatiles (both synthetic and natural ones produced by host plants) by sunlight and wind in the canopy hinders detection by Euglossines, which would explain canopy avoidance [^74^]. Interestingly, these bees also showed the strongest functional response to recovery and stratification. Functional richness was significantly higher in forest understories of less isolated plots, where communities were the most conserved, comprising smaller, fast-flying bees with relatively longer proboscises. The small and filtered subset of species in canopies was variable but tended to contain larger bees more apt to maneuverability, such as those in the genus Eulaema. Unlike stingless bees, these bees seem to adhere to the shade-loving pattern shown by Bawa [^74^], with larger and hairier bees such as Eulaema more prone to successful thermoregulation in more exposed conditions ^87^

While different pollinator groups exhibited their own idiosyncratic recovery trajectories and functional responses to stratification, our study reaffirms the importance of both primeval and successional forests in maintaining animal diversity in disturbed landscapes, as documented for a range of ecosystems [^70,88^]. A high abundance and diversity of pollinators in recently stratified successional forests, as well as unique communities interacting with distinctive sets of plants, underscore the importance of structural complexity in both secondary and primary forests in maintaining and recovering biodiversity [^89^]. However, interaction networks in secondary forests did not resemble those in old growth. Therefore, while passive restoration can be seen as an effective process for fostering pollinator communities in disturbed landscapes, safeguarding a sufficient cover of old primary forests remains necessary to preserve a diverse source of pollinators and plant species in regenerating landscapes. Whenever selective logging cannot be regulated, allowing patches of forest to naturally recover may still significantly increase resilience in disturbed landscapes by providing refuge and non-nutritional resources to pollinators [^90^]. Furthermore, the group-specific interactions between stratification, recovery, and environmental factors highlight the need for comprehensive biodiversity assessments that consider canopies as species-rich sections of the landscape.

## ACKNOWLEDGEMENTS

This work was supported by the Deutsche Forschungsgemeinschaft (DFG) funded Research Unit REASSEMBLY (FOR 5207; sub-projects LE2750/12-1 and KE1742/13-1). We thank the Ministry of Environment of Ecuador for granting research and collection permits through Contrato Marco MAE-DNB-CM-2021-0187, Felicity Newell for collecting the topographic data (Forest in 500 m buffer) for all plots, thank Sebastián Escobar for handling exportation permits, Martin Schaefer (Fundación Jocotoco) and Citlalli Morelos-Juarez (Fundación Tesoro Escondido) for the permission to do research on their reserves, and the staff of both reserves for their immense logistical support: Katrin Krauth, Julio Carbajal, Jender Vélez, Bryan Tamayo, Lady Condoy, Leonardo de la Cruz, Jefferson Tacuri, Vicente Vélez, Alcides Zambrano, Yadira Giler, Patricio Encarnacion, and Adriana Argoti. Daniel Veit (Jena) was instrumental in the construction of the insect funnel traps.

## AUTHOR CONTRIBUTIONS

SDL, GB, AK, and UMD conceptualized the research. SDL, AK and GB acquired and managed the funding. UMD, DB, SVL, and JW performed fieldwork and data collection. UMD, DB, SVL, JW, MP and KF performed sample processing and data curation. CR, GB, UMD, DB, SVL, JW, MP, KF identified insects. UMD performed data analysis and wrote the manuscript draft. All authors contributed critically to the last manuscript draft.

## DATA AVAILABILITY

All the raw data and code scripts used to produce the results of this manuscript will be made publicly available in an online Zenodo repository upon acceptance.

## ADDITIONAL INFORMATION

The authors declare no competing interests.

## REFERENCES

1. Gaston, K. J. Global patterns in biodiversity. Nature 405, 220–227 (2000).

2. Forman, R. T. T. & Godron, M. Landscape Ecology. (Wiley, 1986).

3. Nadkarni, N. M. Diversity of Species and Interactions in the Upper Tree Canopy of Forest Ecosystems. Am Zool 34, 70–78 (1994).

4. Nadkarni, N. M., Merwin, M. C. & Nieder, J. Forest Canopies, Plant Diversity. in Encyclopedia of Biodiversity (Second Edition) (ed. Levin, S. A.) 516–527 (Academic Press, Waltham, 2013). doi:10.1016/B978-0-12-384719-5.00158-1.

5. Shaw, D. C. Vertical organization of canopy biota. in Forest Canopies 73–101 (Lowman, Margaret; Rinker. H. Bruce. Academic Press, 2004).

6. Basset, Y., Hammond, P., Barrios, H., Holloway, J. & Miller, S. Vertical stratification of arthropod assemblages. in 17–27 (2003).

7. Ozanne, C. M. P. et al. Biodiversity Meets the Atmosphere: A Global View of Forest Canopies. Science 301, 183–186 (2003).

8. Baker, H. G., Bawa, K. S., Frankie, G. W. & Opler, P. A. Reproductive biology of plants in tropical forests. in Tropical rainforest ecosystems. 183–215 (F.B. Golley, Elsevier Scientific Publ Co, New Yor, 1983).

9. Ramalho, M. Stingless bees and mass flowering trees in the canopy of Atlantic Forest: a tight relationship. Acta Bot. Bras. 18, 37–47 (2004).

10. Schowalter, T. & Chao, J.-T. Canopy Insect Sampling. in Measuring Arthropod Biodiversity: A Handbook of Sampling Methods (eds. Santos, J. C. & Fernandes, G. W.) 467–493 (Springer International Publishing, Cham, 2021). doi:10.1007/978-3-030-53226-0_18.

11. Asner, G. P. et al. Condition and fate of logged forests in the Brazilian Amazon. Proceedings of the National Academy of Sciences 103, 12947–12950 (2006).

12. Xu, H. et al. Partial recovery of a tropical rain forest a half-century after clear-cut and selective logging. Journal of Applied Ecology 52, 1044–1052 (2015).

13. Myers, N., Mittermeier, R. A., Mittermeier, C. G., da Fonseca, G. A. B. & Kent, J. Biodiversity hotspots for conservation priorities. Nature 403, 853–858 (2000).

14. Silva, I., Rocha, R., Lopez-Baucells, A., Farneda, F. & Meyer, C. Effects of Forest Fragmentation on the Vertical Stratification of Neotropical Bats. Diversity 12, 67 (2020).

15. Whitworth, A. et al. Past Human Disturbance Effects upon Biodiversity are Greatest in the Canopy; A Case Study on Rainforest Butterflies. PLOS ONE 11, e0150520 (2016).

16. López-Cubillos, S., McDonald-Madden, E., Mayfield, M. M. & Runting, R. K. Optimal restoration for pollination services increases forest cover while doubling agricultural profits. PLOS Biology 21, e3002107 (2023).

17. Nakamura, A. et al. Forests and Their Canopies: Achievements and Horizons in Canopy Science. Trends in Ecology & Evolution 32, 438–451 (2017).

18. Barker, M. & Pinard, M. Forest canopy research: Sampling problems, and some solutions. Plant Ecology 153, 23–38 (2001).

19. Brehm, G. Contrasting patterns of vertical stratification in two moth families in a Costa Rican lowland rain forest. Basic and Applied Ecology 8, 44–54 (2007).

20. Roubik, D. W. Ecology and Natural History of Tropical Bees. (Cambridge University Press, 1992).

21. Stangler, E. S., Hanson, P. E. & Steffan-Dewenter, I. Vertical diversity patterns and biotic interactions of trap-nesting bees along a fragmentation gradient of small secondary rainforest remnants. Apidologie 47, 527–538 (2016).

22. Allen, G. & Davies, R. G. Canopy sampling reveals hidden potential value of woodland trees for wild bee assemblages. Insect Conservation and Diversity 16, 33–46 (2023).

23. Ashton, L. A. et al. Vertical stratification of moths across elevation and latitude. Journal of Biogeography 43, 59–69 (2016).

24. Roubik, D. W. Tropical pollinators in the canopy and understory: Field data and theory for stratum “preferences”. J Insect Behav 6, 659–673 (1993).

25. Smith, S. M., Islam, N. & Bellocq, M. I. Effects of single-tree selection harvesting on hymenopteran and saproxylic insect assemblages in the canopy and understory of northern temperate forests. Journal of Forestry Research 23, 275–284 (2012).

26. Kirmse, S. & Chaboo, C. S. Flowers are essential to maintain high beetle diversity (Coleoptera) in a Neotropical rainforest canopy. Journal of Natural History 54, 1661– 1696 (2020).

27. Escobar, S. et al. Reassembly of a tropical rainforest ecosystem: A new chronosequence in the Ecuadorian Chocó tested with the recovery of tree attributes. 2024.03.21.586145 Preprint at 10.1101/2024.03.21.586145 (2024).

28. Brehm, G. A new LED lamp for the collection of nocturnal Lepidoptera and a spectral comparison of light-trapping lamps. Nota Lepidopterologica 40, 87–108 (2017).

29. Renner, S. S. & Feil, J. P. Pollinators of Tropical Dioecious Angiosperms. American Journal of Botany 80, 1100–1107 (1993).

30. Diniz, U. M. et al. The neglected pollinators: settling moths are keystone floral visitors essential to network connectivity and tropical forest recovery. Proceedings of the Royal Society B: Biological Sciences early view, (2025).

31. Rentería, E. & Brehm, G. Is Blue the Most Attractive Color for Bees? Exploring the Attractiveness of Colors in Vane Traps. J Insect Behav 38, 28 (2025).

32. Michener, C. D. The Bees of the World. (Johns Hopkins University Press, 2007). doi:10.56021/9780801885730.

33. Brehm, G., et al. Illustrated catalogue and preliminary phylogeny of 330 species of Arctiinae moth species from the Chocó rainforest in NW Ecuador: most species are undescribed. https://ecoevorxiv.org/repository/view/8853/ (2025).

34. Bonilla-Gómez, M. A. & Nates-Parra, G. Abejas Euglosinas De Colombia (hymenoptera: Apidae) I. Claves Ilustradas. Caldasia 17, 149–172 (1992).

35. Roubik, D. W. Stingless Bees: A guide to Panamanian and Mesoamerican species and their nests (Hymenoptera: Apidae: Meliponinae). in Insects of Panama and Mesoamerica: Selected Studies (eds. Quintero, D. & Aiello, A.) 0 (Oxford University Press, 1992). doi:10.1093/oso/9780198540182.003.0031.

36. Bailey, R. I., Molleman, F., Vasseur, C., Woas, S. & Prinzing, A. Large body size constrains dispersal assembly of communities even across short distances. Sci Rep 8, 10911 (2018).

37. Cane, J. H. Estimation of Bee Size Using Intertegular Span (Apoidea). Journal of the Kansas Entomological Society 60, 145–147 (1987).

38. Johnson, S. D. et al. The long and the short of it: a global analysis of hawkmoth pollination niches and interaction networks. Functional Ecology 31, 101–115 (2017).

39. Kunte, K. Allometry and functional constraints on proboscis lengths in butterflies. Functional Ecology 21, 982–987 (2007).

40. Bhat, S. S., Zhao, J., Sheridan, J., Hourigan, K. & Thompson, M. C. Aspect ratio studies on insect wings. Physics of Fluids 31, 121301 (2019).

41. Collins, T. J. ImageJ for Microscopy. BioTechniques 43, S25–S30 (2007).

42. Jaimes Nino, L. M., Mörtter, R. & Brehm, G. Diversity and trait patterns of moths at the edge of an Amazonian rainforest. J Insect Conserv 23, 751–763 (2019).

43. Badejo, O., Skaldina, O., Gilev, A. & Sorvari, J. Benefits of insect colours: a review from social insect studies. Oecologia 194, 27–40 (2020).

44. Sickel, W. et al. Increased efficiency in identifying mixed pollen samples by meta-barcoding with a dual-indexing approach. BMC Ecology 15, 20 (2015).

45. Campos, M. G. et al. Standard methods for pollen research. Journal of Apicultural Research 60, 1–109 (2021).

46. Keller, A. et al. Evaluating multiplexed next-generation sequencing as a method in palynology for mixed pollen samples. Plant Biology 17, 558–566 (2015).

47. Leonhardt, S. D., Peters, B. & Keller, A. Do amino and fatty acid profiles of pollen provisions correlate with bacterial microbiomes in the mason bee Osmia bicornis? Philosophical Transactions of the Royal Society B: Biological Sciences 377, 20210171 (2022).

48. Rognes, T., Flouri, T., Nichols, B., Quince, C. & Mahé, F. VSEARCH: a versatile open source tool for metagenomics. PeerJ 4, e2584 (2016).

49. Keller, A. et al. BCdatabaser: on-the-fly reference database creation for (meta-)barcoding. Bioinformatics 36, 2630–2631 (2020).

50. Quaresma, A. et al. Semi-automated sequence curation for reliable reference datasets in ITS2 vascular plant DNA (meta-)barcoding. Sci Data 11, 129 (2024).

51. Edgar, R. C. SINTAX: a simple non-Bayesian taxonomy classifier for 16S and ITS sequences. 074161 Preprint at 10.1101/074161 (2016).

52. McMurdie, P. J. & Holmes, S. phyloseq: An R Package for Reproducible Interactive Analysis and Graphics of Microbiome Census Data. PLOS ONE 8, e61217 (2013).

53. Core Team, R. C. T. R: A Language and Environment for Statistical Computing. R Foundation for Statistical Computing. R Foundation for Statistical Computing. https://www.r-project.org/ (2025).

54. Oksanen, J. et al. vegan: Community Ecology Package. (2025).

55. Dormann, C. F., Gruber, B. & Fründ, J. Introducing the bipartite Package: Analysing Ecological Networks. R News 2, (2008).

56. Laliberté, E. & Legendre, P. A distance-based framework for measuring functional diversity from multiple traits. Ecology 91, 299–305 (2010).

57. Baselga, A. & Orme, C. D. L. betapart: an R package for the study of beta diversity. Methods in Ecology and Evolution 3, 808–812 (2012).

58. Magurran, A. E. Ecological Diversity and Its Measurement. (Springer Science & Business Media, 2013).

59. Villéger, S., Mason, N. W. H. & Mouillot, D. New Multidimensional Functional Diversity Indices for a Multifaceted Framework in Functional Ecology. Ecology 89, 2290–2301 (2008).

60. Blüthgen, N., Menzel, F. & Blüthgen, N. Measuring specialization in species interaction networks. BMC Ecol 6, 9 (2006).

61. Rader, R. & Krockenberger, A. Does resource availability govern vertical stratification of small mammals in an Australian lowland tropical rainforest? Wildl. Res. 33, 571–576 (2006).

62. Fermon, H., Waltert, M., Vane-Wright, R. I. & Mühlenberg, M. Forest use and vertical stratification in fruit-feeding butterflies of Sulawesi, Indonesia: impacts for conservation. Biodivers Conserv 14, 333–350 (2005).

63. Basham, E. W., Baecher, J. A., Klinges, D. H. & Scheffers, B. R. Vertical stratification patterns of tropical forest vertebrates: a meta-analysis. Biological Reviews 98, 99–114 (2023).

64. Kalko, E. K. V. & Handley, C. O. Neotropical Bats in the Canopy: Diversity, Community Structure, and Implications for Conservation. Plant Ecology 153, 319–333 (2001).

65. Sobek, S., Tscharntke, T., Scherber, C., Schiele, S. & Steffan-Dewenter, I. Canopy vs. understory: Does tree diversity affect bee and wasp communities and their natural enemies across forest strata? Forest Ecology and Management 258, 609–615 (2009).

66. Ulyshen, M. D., Soon, V. & Hanula, J. L. On the vertical distribution of bees in a temperate deciduous forest. Insect Conservation and Diversity 3, 222–228 (2010).

67. Schulze, C. H., Linsenmair, K. E. & Fiedler, K. Understorey versus canopy: patterns of vertical stratification and diversity among Lepidoptera in a Bornean rain forest. in Tropical Forest Canopies: Ecology and Management: Proceedings of ESF Conference, Oxford University, 12–16 December 1998 (eds. Linsenmair, K. E., Davis, A. J., Fiala, B. & Speight, M. R.) 133–152 (Springer Netherlands, Dordrecht, 2001). doi:10.1007/978-94-017-3606-0_11.

68. Calaça, P., de Freitas, L. D. & Schlindwein, C. Strongly unbalanced gender attractiveness in a dioecious mass flowering tropical tree pollinated by stingless bees. Plant Biology 24, 473–481 (2022).

69. Chazdon, R. Chance and Determinism in Tropical Forest Succession. in (2008).

70. Taki, H. et al. Evaluation of secondary forests as alternative habitats to primary forests for flower-visiting insects. J Insect Conserv 17, 549–556 (2013).

71. Fiedler, K., Hilt, N., Brehm, G. & Schulze, C. H. Moths at tropical forest margins — how mega-diverse insect assemblages respond to forest disturbance and recovery. in Stability of Tropical Rainforest Margins: Linking Ecological, Economic and Social Constraints of Land Use and Conservation (eds. Tscharntke, T., Leuschner, C., Zeller, M., Guhardja, E. & Bidin, A.) 37–58 (Springer, Berlin, Heidelberg, 2007). doi:10.1007/978-3-540-30290-2_3.

72. Ulyshen, M., Urban-Mead, K. R., Dorey, J. B. & Rivers, J. W. Forests are critically important to global pollinator diversity and enhance pollination in adjacent crops. Biological Reviews 98, 1118–1141 (2023).

73. Bihn, J. H., Verhaagh, M., Brändle, M. & Brandl, R. Do secondary forests act as refuges for old growth forest animals? Recovery of ant diversity in the Atlantic forest of Brazil. Biological Conservation 141, 733–743 (2008).

74. Bawa, K. S. Plant-Pollinator Interactions in Tropical Rain Forests. Annual Review of Ecology, Evolution, and Systematics 21, 399–422 (1990).

75. Wardhaugh, C. W., Stork, N. E. & Edwards, W. Canopy invertebrate community composition on rainforest trees: Different microhabitats support very different invertebrate communities. Austral Ecology 39, 367–377 (2014).

76. Xing, S. et al. Ecological patterns and processes in the vertical dimension of terrestrial ecosystems. Journal of Animal Ecology 92, 538–551 (2023).

77. González del Pliego, P., et al. Thermally buffered microhabitats recovery in tropical secondary forests following land abandonment. Biological Conservation 201, 385–395 (2016).

78. TOLEDO-HERNÁNDEZ, E. et al. The stingless bees (Hymenoptera: Apidae: Meliponini): a review of the current threats to their survival. Apidologie 53, 8 (2022).

79. Samejima, H., Marzuki, M., Nagamitsu, T. & Nakasizuka, T. The effects of human disturbance on a stingless bee community in a tropical rainforest. Biological Conservation 120, 577–587 (2004).

80. Chui, S. X., Keller, A. & Leonhardt, S. D. Functional resin use in solitary bees. Ecological Entomology 47, 115–136 (2022).

81. Grüter, C. & Hayes, L. Sociality is a key driver of foraging ranges in bees. Current Biology 32, 5390–5397.e3 (2022).

82. Gonzalez, V. H. et al. Low heat tolerance and high desiccation resistance in nocturnal bees and the implications for nocturnal pollination under climate change. Sci Rep 13, 22320 (2023).

83. Ramos, Y. J. A., Romero, E. & Gálvez, D. Vertical stratification in orchid bees (Apidae: Euglossini)?: a meta-analysis. Apidologie 53, 26 (2022).

84. Eltz, T., Whitten, W. M., Roubik, D. W. & Linsenmair, K. E. Fragrance Collection, Storage, and Accumulation by Individual Male Orchid Bees. J Chem Ecol 25, 157–176 (1999).

85. Rasmussen, C. Diversity and abundance of orchid bees (Hymenoptera: Apidae, Euglossini) in a tropical rainforest succession. Neotrop. entomol. 38, 66–73 (2009).

86. Feitosa Ribeiro, C., et al. Vertical distribution of orchid bees (Hymenoptera: Apidae: Euglossini) in an Amazon forest fragment. Journal of Apicultural Research 63, 1058–1066 (2024).

87. May, M. L. & Casey, T. M. Thermoregulation and Heat Exchange in Euglossine Bees. Physiological Zoology 56, 541–551 (1983).

88. Ulyshen, M. D., Ballare, K. M., Fettig, C. J., Rivers, J. W. & Runyon, J. B. The Value of Forests to Pollinating Insects Varies with Forest Structure, Composition, and Age. Curr. For. Rep. 10, 322–336 (2024).

89. Hilmers, T. et al. Biodiversity along temperate forest succession. Journal of Applied Ecology 55, 2756–2766 (2018).

90. Meli, P. et al. A global review of past land use, climate, and active vs. passive restoration effects on forest recovery. PLOS ONE 12, e0171368 (2017).

