## Supplementary Material for "Canopies drive the reassembly of pollinator communities and interaction networks along tropical forest succession"

**Box S1**. Bow-and-arrow troubleshooting – common issues and quick fixes

**Issue 1. The line or knot snaps during shooting and the arrow is lost.** Provided that a quality knot has been used, this issue is usually caused by a loose line that is not rolled up tightly enough onto the spool of the fishing reel. Loose lines create small segments in which the line tangles in itself and thus does not unroll properly, creating enough resistance to cause it or the knot to snap when the arrow is shot. If this happens, the line roll on the spool will be clearly deformed, and the line must be entirely unrolled from the spool and re-rolled with enough tension. Other factors that might contribute to the issue are a tight drag adjustment (must be completely loosened up before the shot) and a wrong angle of the reel in relation to the arrow (see Fig. 1, A).

**Issue 2. The arrow passes below the branch.** Normally caused by shooting at an incorrect angle. The arrow must be aimed slightly above the branch. If there is no vegetation clutter above the branch, the arrow can also be shot widely above it. Heavy fishing lines (e.g., nylon) or an improperly rolled-up line on the spool (see above) might create resistance and pull, respectively, decelerating the arrow and contributing to the issue.

**Issue 3. The arrow passes above the branch but does not come down from the canopy.** The most common issue, which occurs even when the arrow is shot correctly. This is caused by the arrow getting tangled in branched and vegetation near or behind the branch. This problem can usually be solved by selecting a fitting, clutter-free branch, as well as one with little vegetation behind it that might stop the trajectory of the arrow. If this problem is recurrent, switching trees is advised. Whenever an arrow gets tangled, a repetitive up-and-down movement with the reel might help to free the arrow.

**Issue 4. The arrow goes over the branch and down from the canopy, but stops midway to the ground.**

Two major causes, which may act synergistically: 1 – Misplaced weight balance. Either the arrow is too light and cannot counterbalance the weight of the line behind it, or the line is too heavy and/or elastic (e.g., some nylon fishing lines), or a combination of both. Light, non-elastic, and sturdy materials are recommended for lines, such as polyethylene. Carbon fiber or glass fiber arrows are ideal; additional weight can be added to arrows by adding pieces of duct tape on the shaft. 2 – The line has many contact points in the canopy, passing through several branches or leaves, creating several resistance points. Shots must ideally only go through one single resistance point. If a single branch cannot be isolated after multiple shot attempts, consider switching trees.

**Issue 5. The arrow passes over the branch and reaches the ground, but is nowhere to be found.** Arrows can end several dozen meters away from the target tree, depending on the shooting angle, and can blend into the vegetation. Shooting from a very diagonal angle (e.g., 45°) in relation to the branch may cause arrows to end up farther away. Shooting at an angle close to 90° (but never entirely vertical) will increase the chances that the arrow ends up near the targeted tree. Bright-colored markings on the arrow greatly reduce search times.

**Issue 6. After tying one extremity of the line to the rope, pulling the line back with the reel is hard to impossible.** Normally caused by the rope knot encountering resistance against the branches in the canopy. A repetitive, up-and-down movement with the reel is normally enough to free the rope. If the rope reaches a point where it’s impossible to pull, it has likely encountered an intersection between two large branches and the knot is too large to pass through. Avoid large, bulky knots and always aim at horizontal branches for setting the rope instead of a V-shaped intersection.

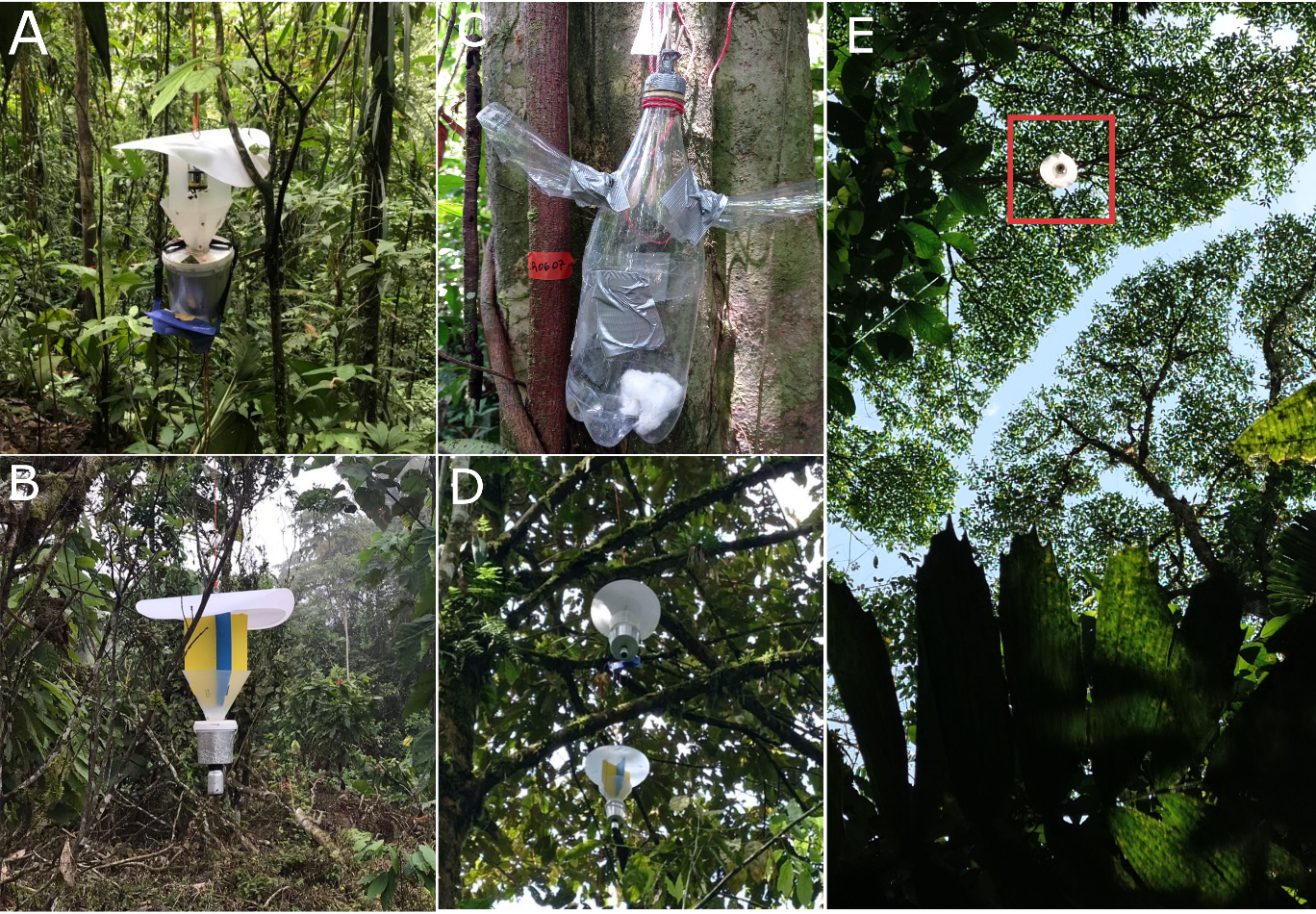

**Figure S1.** Traps used in the study and their location in the canopy. A – The white vane trap equipped with a LepiLED mixed UV light, set in the understory. The blue dry bag contains a power bank and a timer. B – The colored vane trap used to capture diurnal bees. C – A fragrance trap used to trap male orchid bees. The trap contains two openings to a chamber containing the attractive compound. D – A close-up of a trap setup in the pulley rope. E – A view of the canopy traps from the understory (red square).

**Box S2.** Description of the traps used to capture pollinators in the study site

**Light vane traps (Fig. S1A):** These traps target nocturnal insects, and in our study captured moths and nocturnal bees. They were modified funnel traps consisting of a roof, three white vanes, and a funnel of white polypropylene, equipped with a LepiLED mini Switch light (1) and connected to a power bank battery. These traps have proven to be highly efficient in capturing a representative diversity of moths in comparison to common active methods (e.g., light sheets) (2). We used the mixed light setting (UV and visible light spectrums). The funnel traps were attached to a collection chamber, a 5 L bucket (wrapped in aluminum foil to avoid overheating) attached to a 100 mL bottle containing chloroform. Chloroform vapors were used as a fast-killing agent; vapors were slowly released into the collection bucket through a cotton rope. The traps were equipped with a timer fed by a portable power bank and set to start light emission at 1800h and stop at 0600h. All trapped insects were collected the following day.

**Color vane traps (Fig. S1B)**: These traps target diurnal bees. The traps were the same as those used by (3) and consisted of a similar setup of the light vane traps above, but with two colored vanes (yellow and blue) attached transversally, forming an alternating color pattern designed to visually attract bees. Insects were collected in a connected 1 L plastic bucket, which had the same setup and killing agent as described above for the light traps. Traps were set during the morning and collected 24h later.

**Fragrance traps (Fig. S2C):** These traps target fragrance-collecting male orchid bees (Apidae: Euglossini), a dominant pollinator group in the Neotropics (3). There is no systematic and widespread method to collect females passively. These traps consisted of 2L PET bottles containing two lateral entrance holes (approximately 3 cm) equipped with plastic funnels as entrances, similar to those in (4). Inside, a piece of cotton soaked in household insecticide (Raid, USA) was used as a fast-killing agent. On each plot, we placed four fragrance traps at the canopy and understory, each containing one of the following compounds: 1,8-cineole, eugenol, methyl-salicylate, and skatol (Carl Roth GmbH, Germany).

**REFERENCES**

1.	Brehm G. A new LED lamp for the collection of nocturnal Lepidoptera and a spectral comparison of light-trapping lamps. Nota Lepidopterol. 2017 Apr 24;40(1):87–108.

2.	Singh RP, Böttger D, Brehm G. Moth light traps perform better with vanes: A comparison of different designs. J Appl Entomol. 2022;146(10):1343–52.

3.	Michener CD. The Bees of the World [Internet]. Johns Hopkins University Press; 2007 [cited 2024 Feb 15]. Available from: https://www.press.jhu.edu/books/title/9040/bees-world

4.	Ferreira RP, Martins C, Dutra MC, Mentone CB, Antonini Y. Old Fragments of Forest Inside an Urban Area Are Able to Keep Orchid Bee (Hymenoptera: Apidae: Euglossini) Assemblages? The Case of a Brazilian Historical City. Neotrop Entomol. 2013 Oct 1;42(5):466–73.
